# Integrative analysis of 5-methyl- and 5-hydroxymethylcytosine indicates a role for 5-hydroxymethylcytosine as a repressive epigenetic mark

**DOI:** 10.1101/318808

**Authors:** Christopher E. Schlosberg, John R. Edwards

## Abstract

Since the discovery of 5-hydroxymethylcytosine (5hmC) as a prominent DNA base modification found in mammalian genomes, an emergent question has been what role this mark plays in gene regulation. 5hmC is hypothesized to function as an intermediate in the demethylation of 5-methylcytosine (5mC) and also in reactivation of silenced regulatory elements, including promoters and enhancers. Further, weak positive correlations have been observed between gene body 5hmC and gene expression. We previously demonstrated that ME-Class, which uses a high-resolution model of whole-genome bisulfite sequencing data, is an effective tool to understand relationships between 5mC and expression. In this work, we present ME-Class2, a machine-learning based tool to perform integrative 5mCG, 5hmCG and expression analysis. Using ME-Class2 we analyze whole-genome single-base resolution 5mC and 5hmC datasets from 20 primary tissue and cell samples to uncover relationships between 5hmC and expression. The addition of 5hmC improves model performance for tissues with high-levels of 5hmC such as the brain. Our analysis further indicates that conversion of 5mC to 5hmC within 2kb of the transcription start site associates with distinct functions depending on the summed level of 5mC + 5hmC. Unchanged levels of 5mC + 5hmC (i.e. conversion from 5mC to stable 5hmC) associate with repression. Meanwhile, decreases in 5mC + 5hmC (i.e. 5hmC-mediated demethylation) associate with gene activation. As more large-scale, genome-wide, differential DNA methylation studies become available, tools such as ME-Class2 will prove invaluable to interpret epigenomic data and guide mechanistic studies into the function of 5hmC.

## INTRODUCTION

In mammalian genomes, cytosines are frequently covalently modified at the 5-position with methyl-, hydroxymethyl-, formyl-, and carboxy-groups. The initial modification occurs by addition of a methyl-group to the 5-position of the cytosine (5-methylcytosine, 5mC) by a DNA methyltransferase (Dnmt). The subsequent modifications are then formed through successive oxidation of 5mC by the Ten-eleven translocation (Tet) family of enzymes. While 5mC occurs at nearly 70% of all CG dinucleotides (CpG) in the genome in all tissues, 5-hydroxymethylcytosine (5hmC) appears to be primarily limited to embryonic stem cells, neurons, liver, breast, testis, and placenta tissues, occurring at 2-17% of CpGs depending on the tissue type (1–3). Meanwhile 5hmC’s oxidized derivatives, 5-formylcytosine (5fC) and 5- carboxycytosine (5caC), are only found at very low levels, 10-1000 fold less than 5hmC (1). It is still unclear whether 5fC and 5caC are short-lived intermediates (2) or whether they have an independent biological function *in vivo* (3, 4). Of these marks, 5mC has been the most studied and is a well-established player in maintaining inactivation of the silenced X chromosome, mono-allelic gene expression at imprinted loci, and silencing retrotransposons (5). Abnormal patterns of 5mC are also linked to transcriptional dysregulation in cancer.

At the biochemical level, 5hmC likely plays a role in demethylation through both passive and active mechanisms. While Dnmt1 is responsible for copying and propagating 5mC during cell division, no similar mechanism has yet been discovered for 5hmC. Notably, 5hmC, 5fC, and 5caC are found at their highest levels in post-mitotic cells and are passively diluted during cell division. Active demethylation occurs through conversion of 5mC to 5caC via 5hmC and 5fC intermediates. 5caC is then converted to unmethylated cytosine through base excision repair or decarboxylation (6). In support of 5hmC’s role as an intermediate in demethylation *in vivo*, Tet2-/- mouse brains exhibit low level gains in methylation (7).

What is known about the biological roles of 5hmC and the Tet family of enzymes (Tet1, Tet2, Tet3) are mostly supported through a combination of genetic and correlative studies. Genomic analyses show that while the majority of the genome is marked by 5mC except for CpG islands (CGIs), gene promoters, and enhancers, 5hmC is enriched at enhancers, gene bodies, and CGI shores. Further, 5hmC is depleted from CGIs in ES cells and neurons and depleted from intergenic regions in ES cells (4–6). 5hmC starts at low levels in the developing brain, but accumulates in the adult brain (8). Tet1-/- mice do not show impaired differentiation (1, 9) and have normal brain morphology (10). However, they do show impairments in synaptic plasticity and memory extinction (10). Tet1-, Tet2-, Tet3-triple KO mice displayed impaired differentiation and embryonic development, and significant promoter hypermethylation (9). These data suggest compensatory effects between Tet family members. 5hmC may also play a role in enhancer regulation, as Tet2 deletion causes an increase in enhancer 5mC levels and reduced enhancer activity (11).

There has been some evidence that 5hmC can play a regulatory role independent of its role as an intermediate in demethylation in post-mitotic neurons and ES cells. For example, MeCP2 displays reduced affinity for hmCG compared to mCG, and therefore conversion of mCG to stable hmCG in the neuronal genome may lead to loss of functional binding sites for MeCP2 (11–13). However, screens for 5hmC interacting factors have uncovered few 5hmC-specific interactors (4). In general, 5hmC in gene bodies is frequently associated with gene expression; this includes both as a stable mark in neurons, and as an intermediate for demethylation. More recent studies in human liver and lung tissues observed 5hmC as a marker of active transcription associated with H3K4me1 at CpG island shores (12). However, whether 5hmC plays a role in promoter regulation is still unclear, and how 5mC and 5hmC signals in promoters and gene bodies synergize to affect gene silencing has not been studied.

We previously developed ME-Class to model methylation both at the gene promoters and in gene bodies to identify genes with a high probability of association between 5mC and gene expression (13). Here, we extend its functionality to incorporate 5hmC to systematically interrogate how changes in 5mC and 5hmC associate with gene expression. Our results indicate that models that include both 5mC and 5hmC out-perform 5mC only models, but only in tissues or cells (such as neuronal tissues) that have high levels of 5hmC. Further, our results indicate that 5hmC associates with gene activation when it is involved in demethylation and with gene repression when it is stably present at and around the promoter of a gene.

## MATERIALS AND METHODS

### WGBS, TAB-seq, oxBS-seq, and RNA-seq data

Mapped sequence reads for whole genome bisulfite sequencing (WGBS), Tet-assisted bisulfite sequencing (TAB-seq), and RNA-seq in liver and lung tumor and matched normal samples were obtained from Li et al. (12) and from dendritic cells from Pacis et al. (14). WGBS, oxidative bisulfite sequencing (oxBS-seq), and RNA-seq from fetal and 6 week mouse brain samples were obtained from Lister et al. (7) and granule cells from Mellen et al. (15). WGBS and TAB-seq for human cortex are from Wen et al. (16). Corresponding RNA-seq data are from Brawand et al. (17) as used by Wen et al. (16).

### Estimation of 5mC and 5hmC levels

5mC and 5hmC levels were estimated using maximum likelihood methylation levels (MLML) from either TAB-seq or oxBS-seq (18). MLML provides a simultaneous maximum likelihood based on binomial estimates of 5hmC and 5mC. We used MLML with a significance level of α=0.05 for the binomial test at each CpG site and an expectation maximization convergence threshold of 1e-10. Counts of individual CpGs with estimated 5hmC and 5mC in all samples can be found in Supplementary Table S1.

### Differential Expression from RNA-seq

RNA-seq data from human liver, lung, and cortex samples were mapped to hg19 using HISAT2 (19). We used featureCounts to estimate feature counts over RefSeq reads (20). Differentially expressed genes were defined as abs[fold change] >=2 after applying a floor of cpm=1. To create a standardized gene set with high quality methylation data, we excluded genes with ambiguous or incomplete transcription start site (TSS) annotations, genes shorter than 5kb, genes with <40 CpGs assayed within +/-5kb of the TSS, genes where, for all CpGs within +/-5kb of the TSS, the change in methylation (mCG/CG) was less than 0.2, and genes with alternative promoters. These filters were used to exclude non-coding genes, pseudogenes, genes shorter than the interpolation boundary (see HRPS model description below), genes with low numbers of CpGs (to reduce bias caused by error in individual CpG measurements), and genes with no methylation changes at their respective promoters. We only included RefSeq genes with cdsStartStat and cdsEndStat flags marked as ‘cmpl’ according to the UCSC Table Browser. For any RefSeq genes with multiple RefSeq IDs corresponding to the same TSS location, we used a single RefSeq ID with the lowest accession number and excluded the remainder. This is a conservative method to simplify the annotations of genes with alternative promoter annotations. A full summary of differentially expressed filtered gene counts can be found in Supplementary Table S2.

### 5hmC incorporation in ME-Class

MLML produces an estimate of 5mC and 5hmC for each CpG site. ME-Class high-resolution promoter signature (HRPS), region of interest (ROI), and whole-scale gene (WSG) models described in Schlosberg et al. (13) were extended to add 5hmCG features (Fig. 1a,b). For the HRPS model, 5mCG and 5hmCG data were independently interpolated using PCHIP interpolation and Gaussian smoothing (50bp bandwidth) across the window +/- 5kb relative to each gene’s TSS. Interpolated curves for ∆5hmCG/CG and ∆5mCG/CG (i.e. the difference in 5hmCG and mCG levels between samples) were discretized to create feature vectors for classification using the average methylation in each 20bp segment. Bins for the ROI model were inspired by Lou et al. (21). Differential 5mCG and 5hmCG levels were computed for each bin, which was then used in the resultant feature vector. For the WSG model, 5mCG and 5hmCG data were scaled to a constant length between the TSS and RefSeq annotated transcription end site (TES). Feature vectors were created from 125 bins upstream of the gene, 125 bins downstream of the gene, and 500 bins from the area between the TSS and TES. Differential 5mCG and 5hmCG levels were both computed for the entire set of bins and then combined to form the final feature vector. ∆5mCG/CG corresponds to the 5mC feature vector and ∆5hmCG/CG corresponds to the 5hmC feature vector. ∆5mCG/CG & ∆5hmCG/CG corresponds to concatenating 5mCG and 5hmCG feature vectors together for the classification. ∆5mCG/CG + ∆5hmCG/CG corresponds to summing 5mCG and 5hmCG values before creating the feature vector for classification.

**Figure. 1.**
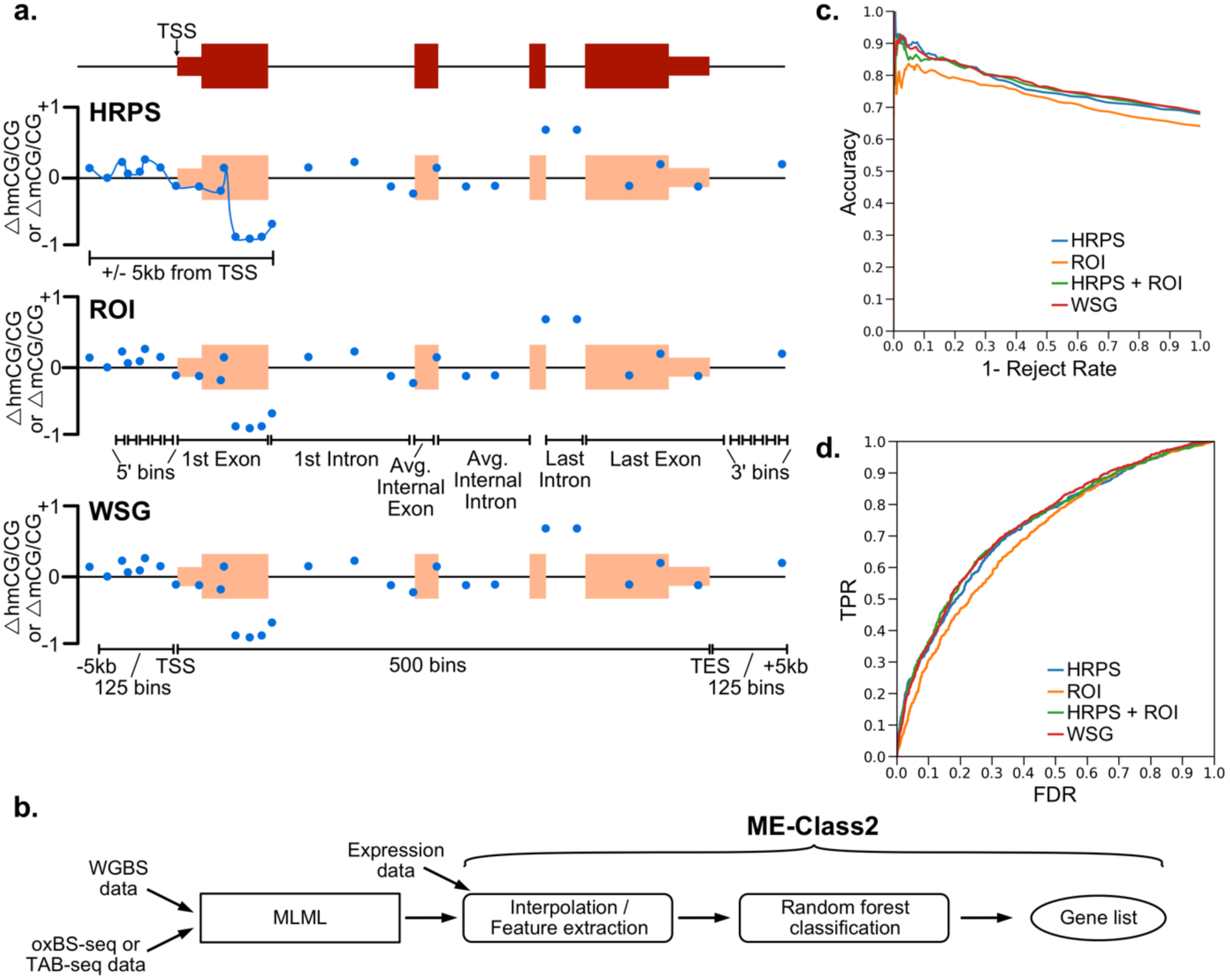
(a) Cartoon example showing different models to encode methylation features for a gene for ME-Class2 analysis. ∆5mCG/CG and ∆5hmCG/CG refer to the differences between two samples. HRPS is high-resolution promoter signature; ROI is region of interest; WSG is whole-scale gene. TSS is transcription start site. Blue dots show example differential methylation (5hmCG or 5mCG). (b) ME-Class2 workflow. (c,d) Performance of different gene models using ME-Class2 5mCG and 5hmCG data from fetal and 6-week mouse brain as evaluated using accuracy versus 1-reject rate (c) and ROC (receiver operating characteristic) curve analysis (d). 1 – reject rate is the fraction of genes with predicted associations between methylation and expression. ROC AUC are HRPS: 0.727, HRPS + ROI: 0.735, WSG: 0.739, ROI: 0.699.

### Evaluation Framework

ME-Class2 uses a random forest classifier which uses feature vectors from 5mCG, 5hmCG or both data to predict the direction of expression change. Random forests were built using 5001 trees. For the fetal to 6-week mouse brain comparison and dendritic cell analysis we used an intra-sample 10-fold cross validation. For the normal liver-lung and normal-tumor comparisons we performed cross-fold validation similar to that in Schlosberg et al. (13). In brief, we hold out each sample one by one for evaluation and then train on the remaining samples. To further minimize over-fitting, all genes from the validation sample are excluded from the samples used for training. For cortex-liver and cortex-lung comparisons, models were trained using 10-fold cross-validation, and all genes used for validation are excluded from the samples used for training.

### Unsupervised Clustering of 5hmCG and 5mC

Unsupervised hierarchical agglomerative clustering (complete linkage) was performed on ∆5mCG/CG and ∆5hmCG/CG in the region [0, +2kb] for the TSS for correctly predicted genes from ME-Class2. Sub-setting our predictions required setting a working threshold for the probability of prediction. Therefore, we set the following range of probabilities of prediction for each experiment based on 90% accuracy at: [0.68, 1.0] fetal-6wk mouse, [0.8, 1.0] normal liver-tumor, [0.7, 1.0] normal-tumor liver and lung. Ranges were set at [0.8-1.0] for cortex-liver and cortex-lung based on 95% accuracy due to the large number of genes accurately predicted for these samples. In the metagene plots of unsupervised results, ∆5mCG/CG+∆5hmCG/CG corresponds to the summation of 5mCG and 5hmCG.

## RESULTS

### Differential 5hmCG at promoter and promoter-proximal regions is more important than gene-body 5hmCG in predicting expression changes

We extended ME-Class to simultaneously incorporate 5mCG and 5hmCG information from high resolution genomic data from WGBS, TAB-seq and oxBS-seq. We used the three best performing models for associating 5mCG and gene expression (13) to understand which model performed the best at the new task of using combined 5mCG and 5hmCG data (Fig. 1a). The first is a high-resolution promoter signature (HRPS) that interpolates a signature around the window +/- 5kb of the TSS for both 5mCG and 5hmCG signals. We previously identified this model as optimal for associating 5mCG and expression changes (13). The second model, which we call “regions of interest” (ROI), bins methylation data upstream of the TSS and across gene features such as first and internal exons and introns (21). We further compared these methods to a whole-scaled gene (WSG) approach, which is based on a scaling method to compare whole gene signals across genes and is commonly used to capture correlations between gene body methylation and expression (8).

We initially benchmarked these models using a set of WGBS and TAB-seq data from fetal and 6-week mouse brains. To evaluate performance, we plot the accuracy versus 1- reject rate for each model. This performance metric allows us to focus on only the genes with the highest quality predictions given some confidence threshold. The underlying premise is that only some genes should have associated DNA methylation and expression changes, not all. We demonstrate good performance to predict gene expression change as measured by both accuracy versus 1-reject rate (Fig. 1c) and ROC analysis (Fig. 1d) for all models. In the HRPS model we predict differential expression in 216 genes with greater than 90% accuracy, which outperforms ROI and WSG models for which we detect a similar number of genes, but at only 83% and 88% accuracy respectively. Both methods that capture the area around the TSS at high resolution (HRPS and WSG) out-perform other methods. Interestingly, models that incorporate features from the gene body (ROI and WSG) do not perform better than those that only model the data around the TSS at high resolution (HRPS). Further, direct addition of gene-body features to the HRPS model (HRPS+ROI) does not increase performance. Random forest feature importance analysis indicates that 5mCG and 5hmCG changes within 2kb and primarily downstream of the TSS into the first intron are the most important regions for successful classification (Fig. 2).

**Figure. 2.**
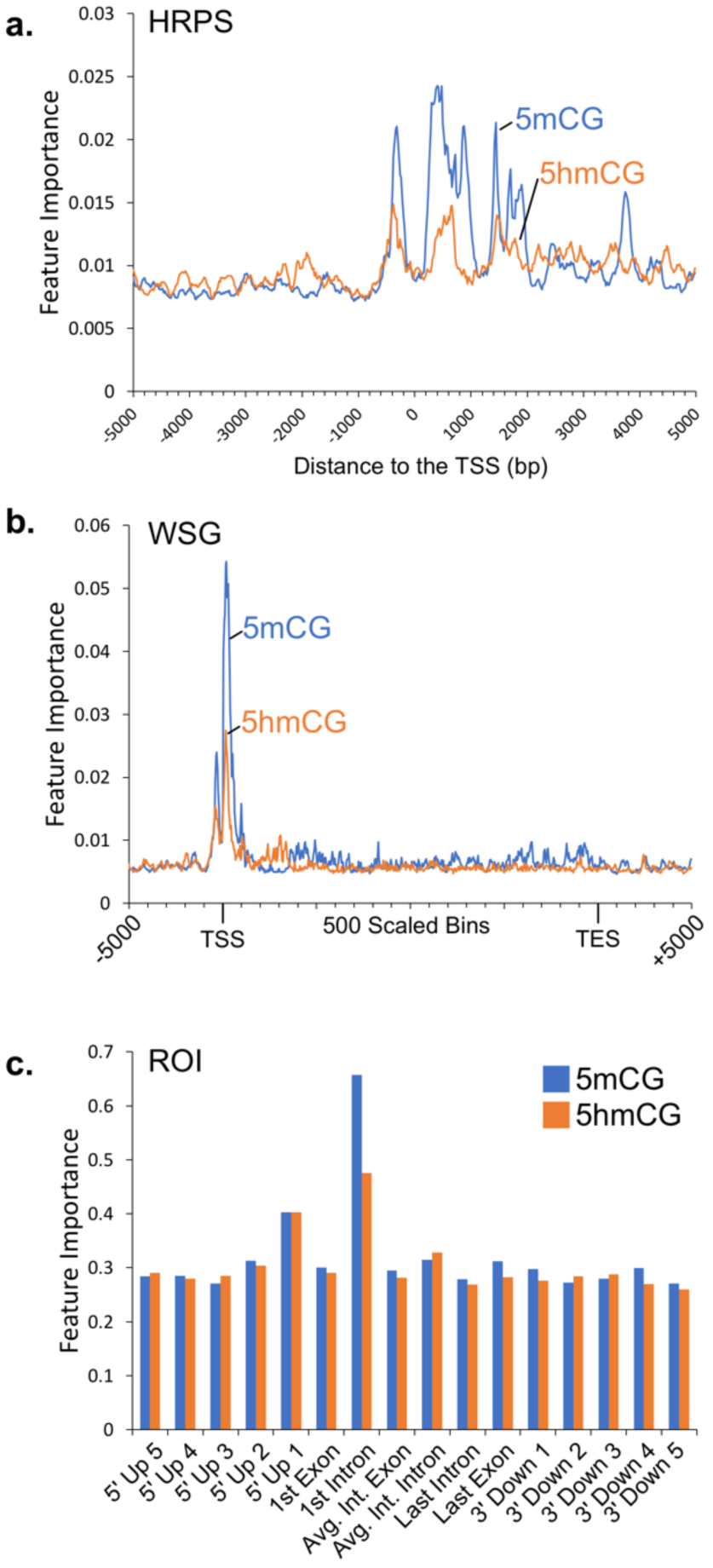
Feature importance for the ME-Class2 random forest classifier for fetal-6wk mouse brain 5mCG and 5hmCG data for (a) HRPS, (b) WSG, and (c) ROI data representations.

### Addition of 5hmCG data improves ME-Class2 performance

We next sought to determine whether models trained using both 5hmCG and 5mcG data outperformed those trained only using 5mCG data. Figure 3a-c shows that including 5mCG and 5hmCG as independent features boosts ME-Class2 performance in the comparison of mouse fetal and 6-week brains and human cortex versus liver and lung using the HRPS model (corresponding ROC curves are in Supplementary Fig. S1). For mouse brain comparisons, the model using 5hmCG and 5mCG data predicted 112 genes at greater than 90% accuracy. Using 5mCG or 5hmCG data alone, the accuracy for a similar number of genes was only 82% and 75% respectively. Similar increases in performance with the inclusion of 5hmCG data were observed for other models including WSG, ROI, and HRPS + ROI (Supplementary Fig. S2). Using the HRPS model, ∆5mCG + ∆5hmC, which is effectively what is measured by only WGBS data in the absence of a 5hmC-specific assay, performed equivalent to ∆5mCG alone (Supplementary Fig. S3).

**Figure. 3.**
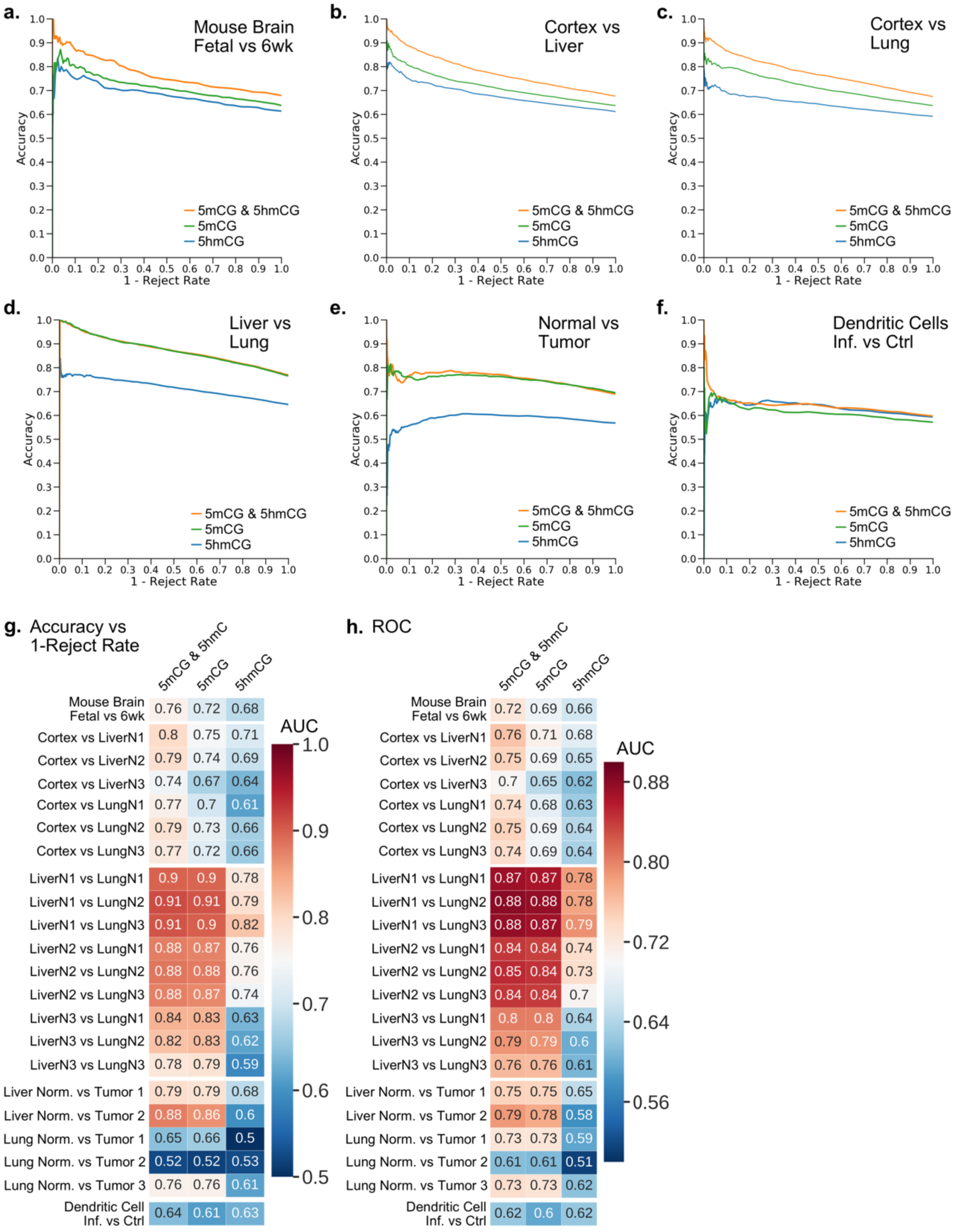
(a-f) ME-Class2 performance (accuracy versus 1-reject rate) for different 5mCG and 5hmCG datasets using the HRPS feature model. The 5mCG and 5mCG & 5hmCG curves directly overlap in panel d. Corresponding detailed ROC (receiver operating characteristic) curves are in Supplementary Fig. S2. (g) Area under the curve (AUC) for the accuracy vs 1-reject rate curves and (h) ROC AUC for each individual sample comparison used in a-f.

We also observe similar performance gains in human cortex vs liver and lung comparisons (Fig. 3). For cortex versus liver, differential expression of an average of 480 genes per sample (493, 493, and 453 for each sample respectively) could be predicted at 90% accuracy using 5hmCG and 5mCG changes, but this accuracy fell to 82% and 77% for 5mCG and 5hmCG alone respectively. Meanwhile for cortex versus lung, differential expression of an average of 278 genes per sample (359, 266, and 209 for each sample respectively) could be predicted at 90% accuracy using 5hmCG and 5mCG changes, but this accuracy fell to 81% and 72% for 5mCG and 5hmCG alone respectively. We also observed an increase in performance comparing bacterially infected and non-infected dendritic cells, although the addition of 5hmCG data only allowed the prediction of differential expression for 16 total genes at greater than 93% accuracy (Fig. 3f, Supplementary Table S3). However, we do not observe such performance gains for all samples. We did not observe any substantial difference between 5mCG only and 5mCG and 5hmCG models in comparisons involving human lung and liver tissues across three individuals (Fig. 3d,g,h), or in normal-tumor comparisons from three lung and two liver tumors (Fig. 3e,g,h). Feature importance analysis of cortex vs liver, cortex vs lung, and infected dendritic cells all support the region within 2-3 kb of the TSS as most important for predicting expression change (Supplementary Fig. S4).

### Predictive methylation signatures in non-brain tissues and tumors are solely dependent on changes in 5mCG

Similar unsupervised clustering of highly predictive liver-lung and cancer-specific genes show why the addition of 5hmCG data did not increase performance in these comparisons (Supplementary Fig. S5). The differential methylation signatures produced from these clusters in each case show that the net ∆5mCG + ∆5hmCG levels closely follow the ∆5mCG levels, with little difference in 5hmCG in all clusters. The observed 5mCG patterns in each cluster resemble those we previously found in other tissues(13) and cancer cell lines (22). This implies that promoter 5mCG, rather than 5hmCG, is primarily associated with gene expression change in cancer.

### ME-Class2 identifies 5hmCG and 5mCG signatures in brain tissues

To better understand why we observed a boost in performance by including 5hmCG in brain and cortex comparisons, we conducted post-hoc unsupervised clustering analysis of identified signatures of 5hmCG and 5mCG that associate with expression change using the mouse fetal and 6-week brain comparison. We observe three distinct classes of differential 5hmCG and 5mCG signatures (Fig. 4a-c). In Figure 4a, we observe increases in both 5mCG and 5hmCG 3’ proximal to the TSS, which associate with a decrease in expression. This contrasts to the signature observed in cluster C2 (Fig. 4b). These genes also decrease in expression; however, while the 5mCG signal decreases 3’ proximal to the TSS, the 5hmCG increases over the same region. There is no substantial change in the net ∆5mCG + ∆5hmCG level, indicating that the primary feature in this cluster is a conversion from 5mCG to stable 5hmCG rather than demethylation. While most of the observed patterns cluster because of changes in 5mC data, observation of the C2 cluster is entirely dependent on the addition of 5hmC data. A third cluster (C3, Fig. 4c) comprises a set of genes that increase in expression and are again characterized by 5mCG decreases and 5hmCG increase 3’ proximal to the TSS. In this case however, the net amount of 5mCG + 5hmCG decreases indicating 5hmCG plays a role as an intermediate toward demethylation. Differential methylation signatures similar to those found in clusters C1 and C3 were also observed in human cortex versus liver and lung comparisons (Supplementary Fig. S6).

**Figure. 4.**
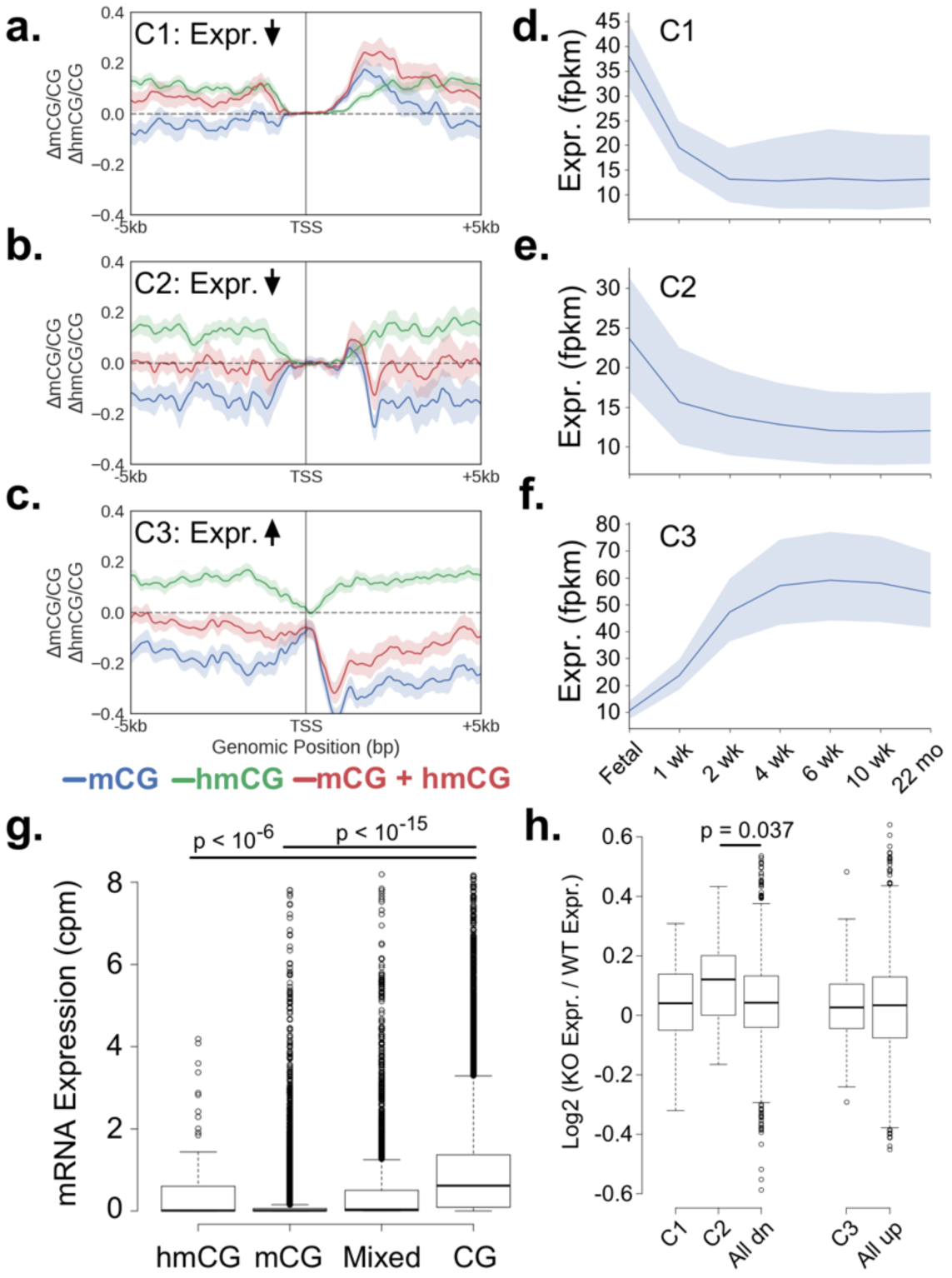
(a-c) Metagene plots for clusters of similar differential methylation signatures (6 wk - fetal) that are predictive of expression in the fetal-6wk mouse brain comparison. Shading indicates the 68% bootstrapped confidence interval. Cluster C1: n=76, C2: n=29, and C3: n=70. (d-f) Average expression of all genes found in C1, C2, and C3 clusters across mouse brain development. Shading indicates the 95% confidence interval. (g) mRNA expression in granule cells of genes whose promoters (defined as [-800bp, +2kb] around the TSS) are greater than 50% marked by mCG, hmCG, a combination of mCG and hmCG, or CG (unmethylated). Outliers have been cropped for clarity. The original plot can be found in Supplementary Fig. S7. (h) Log2 expression changes in cortex from Tet1-/- mice versus cortex from WT mouse. All dn and all up correspond to all down- and up-regulated genes, respectively, in 6wk compared to fetal mouse brain. P-values computed using a Bonferroni adjusted Wilcoxon test.

To better understand whether 5hmCG marked promoters associated with gene repression we examined expression levels of genes in each cluster across mouse development. Cluster C2 genes which are marked by 5mCG alone in the fetal cortex have much lower expression as a whole than genes that gain 5mCG and 5hmCG found in cluster C1 (p < 0.009, Wilcoxon test, Fig. 4d-f). This is in agreement with our finding that 5hmCG within 2kb of the TSS associates with the repression of transcription. To test whether this conclusion would hold true in an alternative dataset, we first used feature importance analysis (Fig. 2a) to identify the region from [-800bp, 2100bp] around the TSS for both 5mCG and 5hmCG signals that contributes the greatest to classification in the fetal-6wk brain comparison. Next, we calculated the average 5mC, 5hmC, and unmodified C content across this region for all genes in granule cells. Agreeing with our hypothesis, genes primarily marked with high levels of either 5mCG (p < 2e-16, Bonferroni adjusted Wilcoxon test) or 5hmCG (p < 2e-7, Bonferroni adjusted Wilcoxon test) are generally not expressed (Fig. 4g).

Lastly to understand whether the conversion or demethylation events are a potential cause of transcriptional change, we examined whether genes from each cluster identified in fetal-6wk comparison above were differentially expressed in Tet1-/- mouse cortex (10). Cluster C1 is characterized by predominantly increased 5mCG levels and thus, as expected, there was no significant difference in the expression of these genes after removal of Tet1. Genes in cluster C2 that were down-regulated in 6-week mouse brain, which had undergone a conversion of 5mCG to 5hmCG (with no net decrease of 5mCG+5hmCG), were found to generally increase in expression in Tet1 -/- mouse cortex relative to WT (p = 0.037, Bonferroni adjusted Wilcoxon test). Surprisingly, there was also no change in expression for genes undergoing Tet-mediated demethylation (cluster C3). This could be because 5hmC-mediated demethylation occurs as a consequence of transcription. In agreement, transcription factor complexes have been implicated to recruit Tet1 leading to 5hmC mediated demethylation mediated by PPARγ in differentiated ES cells (23). However, our analysis has several limitations that could explain the lack of an observed effect. The promoters of selected genes that are differentially expressed in Tet1 -/- cortex and hippocampus were shown to increase in 5mCG levels by only 11-50%, which may be insufficient for many genes to show a change in expression (10). Additionally, we cannot rule out that other Tet members play a compensatory role in the absence of Tet1. In support of the fact that the observed demethylation may activate transcription, a gene found in cluster C3, regulator of G protein signaling RGS14, was shown previously to up-regulate after demethylation of neural progenitors using Dnmt inhibitors(24). In summary, while these data support a role for 5hmCG as a functional repressor, whether demethylation is a cause or consequence of transcriptional silencing or whether there is a context-dependent component is unclear.

### ME-Class2 identifies genes associated with neurodevelopmental disorders and neuronal development

Gene ontology analysis using DAVID (25) revealed genes associated with neurodevelopmental disorders and basic neuronal development in all clusters (Supplementary Table S4). Several of these have been implicated to have differential methylation associated with different disorders including Shank2 in cluster C2 and Nrxn1, Pacsin1, and Grin1 in cluster C3. Shank2, a synaptic protein, has previously been shown to change methylation in the developing human brain and is associated with neurodevelopmental disorders (26). Methylation of Grin1, a component of NMDA receptor complexes, is associated with depression in children (27). Nrxn1 has previously been discovered as having a high ranking meQTL in 110 human hippocampus samples (28). Age-related DNA methylation changes have been found in Nrxn1, which has been implicated in schizophrenia and autism (29). Methylation of PACSIN1 is associated with substance-use risk (30). Importantly, our analysis suggests that 5hmCG may regulate disease-risk genes differently depending on whether it plays a role in repression or demethylation.

## DISCUSSION

We successfully extended ME-Class to predict gene expression classification from both 5hmCG and 5mCG. Feature importance analysis shows that even in tissues with substantial 5hmCG, 5mCG is still the most useful mark for predicting expression changes (Fig. 2, Supplementary Fig. S4). 5hmCG models alone perform very poorly, which demonstrates the importance of considering 5hmC in the context of 5mC to understand potential associations and effects on transcription. Unsupervised analysis revealed a set of down-regulated genes with no net change in 5mCG + 5hmCG levels, but for which 5mCG levels decrease and 5hmCG levels rise. Models using only WGBS data would miss these genes since WGBS only observes the net change in 5mCG + 5hmCG. For other tissues with minimal amounts of 5hmCG it is unlikely that obtaining TAB- or oxBS-seq data will provide more information over what is already found using WGBS (5mCG + 5hmCG).

Our results suggest that the incorporation of 5mCG and 5hmCG marks in the gene body or outside a 2-3kb window from the TSS has little impact on the ability to associate methylation and transcription changes. The direct addition of gene body features based on differential methylation of internal exons and introns using the ROI approach led to no boost in performance. Feature importance analysis (Fig. 2, Supplementary Fig. S4) clearly indicates that for all models, the features within 2-3 kb of the TSS are most essential for prediction and that gene body features greater than 2-3 kb from the TSS are of limited utility. Taken together, this implies either that average gene body 5hmCG plays little functional role in the regulation of transcription, or that gene body information is redundant with that found within 2-3 kb of the promoter. Another alternative is that gene body 5hmCG plays a subtle effect on gene regulation that can only be uncovered with additional training data. For example, these models do not effectively incorporate individual regulatory elements such as enhancers or cryptic promoters found in the gene body that may have context-dependent contributions.

Our results further show that the addition of 5hmCG data has the greatest effect on performance in samples with substantial amounts of 5hmCG such as found in the brain. 5hmCG accumulates in the adult brain as can be observed in Fig. 4a, where 5hmCG increases in most regions around the TSS in 6wk relative to fetal mouse brain. Post-mitotic neurons have high 5hmCG levels and thus these samples benefit the most from inclusion of 5hmCG for predictions. 5hmCG exists at relatively low levels in liver (2.27%-5.68%) and lung (1.94-3.04%) (12) in comparison to mouse brain (17.2%) (7) and human cortex (13%) (16) tissue. 5hmCG is an intermediate cytosine modification which is not replicated during mitosis. Lack of gene expression correlating 5hmCG patterns in normal lung and liver may be because dividing cells in these tissues passively dilute 5hmCG from their genomes. Thus, the scarcity of 5hmCG might explain its lack of predictive ability for expression class change in non-neuronal tissues. In agreement, clustering analysis of 5mCG and 5hmCG signals of predictive genes did not reveal a cluster of 5mCG to 5hmCG conversion as we observed in the model of mouse brain development. Instead, the patterns of differential 5hmCG and 5mCG closely follows that of 5mCG alone across all predictive signatures. However, we cannot rule out that 5hmCG inclusion in tissues with low amounts of 5hmCG might facilitate the identification of a few rare genes regulated by 5hmCG, which cannot be assessed with the limited amount of training data currently available.

Lastly our work points to a potential new mechanism of 5hmCG mediated repression of gene promoters independent of that observed by 5mCG. We identify that conversion of 5mCG to 5hmCG primarily is associated with the downregulation of gene expression and many of these genes are up-regulated upon the removal of Tet1. Since there have been very few proteins identified that specifically bind 5hmC relative to 5mC (4), it is possible that 5hmCG-associated gene silencing could instead be caused by Tet1-recruitment of interacting partners, such as Sin3A and OGT, which have been shown to be involved in Tet1-dependent silencing of LINE-1 (31). It is further unclear at this point why 5hmCG would stabilize in some genes versus other, and complicating matters is that how much active versus passive demethylation occurs via 5hmCG is still a point of contention. It may be that demethylation is the dominant mechanism prior to neurons exiting the cell cycle, while stable 5hmCG occurs after.

ME-Class2 demonstrates that incorporating 5hmCG information is critical for prediction of gene expression changes in samples with high levels of 5hmC such as the brain and neurons. ME-Class2 has identified a class 5mCG/5hmCG patterns that show the conversion from 5mCG to 5hmCG in the 3’ proximal region of the promoter in a model of mouse brain development. We speculate that these patterns of 5mCG and 5hmCG coordinate with additional silencing factors potentially recruited either directly by 5hmCG or by the Tet enzymes in a context-specific manner. As the field continues to collect genome-wide, differential DNA methylation (including both 5mC and 5hmC), tools such as ME-Class2 will prove invaluable for the interpretation of this epigenomic data and will guide mechanistic studies into the integrated function of 5mC and 5hmC in human disease.

## AVAILABILITY

ME-Class2 is an open source collaborative initiative available in the GitHub repository (https://github.com/jredwards417/me-class2)

## ACKNOWLEDGEMENT

We would also like to thank Jerry Fong and Scot Matkovich for critical reading of the manuscript, and Kilian Weinberger and Harrison Gabel for helpful discussions.

## REFERENCES

1. Ito,S., Shen,L., Dai,Q., Wu,S.C., Collins,L.B., Swenberg,J.A., He,C. and Zhang,Y. (2011) Tet proteins can convert 5-methylcytosine to 5-formylcytosine and 5-carboxylcytosine. Science, 333, 1300–1303.

2. Branco,M.R., Ficz,G. and Reik,W. (2012) Uncovering the role of 5-hydroxymethylcytosine in the epigenome. Nat Rev Genet, 13, 7–13.

3. Raiber,E.-A., Murat,P., Chirgadze,D.Y., Beraldi,D., Ben F Luisi and Balasubramanian,S. (2015) 5-Formylcytosine alters the structure of the DNA double helix. Nat Struct Mol Biol, 22, 44–49.

4. Iurlaro,M., Ficz,G., Oxley,D., Raiber,E.-A., Bachman,M., Booth,M.J., Andrews,S., Balasubramanian,S. and Reik,W. (2013) A screen for hydroxymethylcytosine and formylcytosine binding proteins suggests functions in transcription and chromatin regulation. Genome Biol, 14, R119.

5. Edwards,J.R., Yarychkivska,O., Boulard,M. and Bestor,T.H. (2017) DNA methylation and DNA methyltransferases. Epigenetics Chromatin, 10, 23.

6. Kohli,R.M. and Zhang,Y. (2013) TET enzymes, TDG and the dynamics of DNA demethylation. Nature, 502, 472–479.

7. Lister,R., Mukamel,E.A., Nery,J.R., Urich,M., Puddifoot,C.A., Johnson,N.D., Lucero,J., Huang,Y., Dwork,A.J., Schultz,M.D., et al. (2013) Global epigenomic reconfiguration during mammalian brain development. Science, 341, 1237905–1237905.

8. Lister,R., Mukamel,E.A., Nery,J.R., Urich,M., Puddifoot,C.A., Johnson,N.D., Lucero,J., Huang,Y., Dwork,A.J., Schultz,M.D., et al. (2013) Global Epigenomic Reconfiguration During Mammalian Brain Development. Science (New York, N.Y.), 341, 1237905–627.

9. Dawlaty,M.M., Breiling,A., Le,T., Barrasa,M.I., Raddatz,G., Gao,Q., Powell,B.E., Cheng,A.W., Faull,K.F., Lyko,F., et al. (2014) Loss of Tet enzymes compromises proper differentiation of embryonic stem cells. Dev. Cell, 29, 102–111.

10. Rudenko,A., Dawlaty,M.M., Seo,J., Cheng,A.W., Meng,J., Le,T., Faull,K.F., Jaenisch,R. and Tsai,L.-H. (2013) Tet1 is critical for neuronal activity-regulated gene expression and memory extinction. Neuron, 79, 1109–1122.

11. Hon,G.C., Song,C.-X., Du,T., Jin,F., Selvaraj,S., Lee,A.Y., Yen,C.-A., Ye,Z., Mao,S.-Q., Wang,B.-A., et al. (2014) 5mC oxidation by Tet2 modulates enhancer activity and timing of transcriptome reprogramming during differentiation. Mol Cell, 56, 286–297.

12. Li,X., Liu,Y., Salz,T., Hansen,K.D. and Feinberg,A. (2016) Whole-genome analysis of the methylome and hydroxymethylome in normal and malignant lung and liver. Genome Res, 26, 1730–1741.

13. Schlosberg,C.E., Vanderkraats,N.D. and Edwards,J.R. (2017) Modeling complex patterns of differential DNA methylation that associate with gene expression changes. Nucleic Acids Res, 45, 5100–5111.

14. Pacis,A., Tailleux,L., Morin,A.M., Lambourne,J., MacIsaac,J.L., Yotova,V., Dumaine,A., Danckaert,A., Luca,F., Grenier,J.-C., et al. (2015) Bacterial infection remodels the DNA methylation landscape of human dendritic cells. - PubMed - NCBI. Genome Res, 25, 1801–1811.

15. Mellén,M., Ayata,P. and Heintz,N. (2017) 5-hydroxymethylcytosine accumulation in postmitotic neurons results in functional demethylation of expressed genes. Proceedings of the National Academy of Sciences of the United States of America, 114, E7812–E7821.

16. Wen,L., Li,X., Yan,L., Tan,Y., Li,R., Zhao,Y., Wang,Y., Xie,J., Zhang,Y., Song,C., et al. (2014) Whole-genome analysis of 5-hydroxymethylcytosine and 5-methylcytosine at base resolution in the human brain. Genome Biol, 15, R49.

17. Brawand,D., Soumillon,M., Necsulea,A., Julien,P., Csárdi,G., Harrigan,P., Weier,M., Liechti,A., Aximu-Petri,A., Kircher,M., et al. (2011) The evolution of gene expression levels in mammalian organs. Nature, 478, 343–348.

18. Qu,J., Zhou,M., Song,Q., Hong,E.E. and Smith,A.D. (2013) MLML: consistent simultaneous estimates of DNA methylation and hydroxymethylation. Bioinformatics, 29, 2645–2646.

19. Kim,D., Ben Langmead and Salzberg,S.L. (2015) HISAT: a fast spliced aligner with low memory requirements. Nat Methods, 12, 357–360.

20. Liao,Y., Smyth,G.K. and Shi,W. (2014) featureCounts: an efficient general purpose program for assigning sequence reads to genomic features. Bioinformatics, 30, 923–930.

21. Lou,S., Lee,H.-M., Qin,H., Li,J.-W., Gao,Z., Liu,X., Chan,L.L., Kl Lam,V., So,W.-Y., Wang,Y., et al. (2014) Whole-genome bisulfite sequencing of multiple individuals reveals complementary roles of promoter and gene body methylation in transcriptional regulation. Genome Biol, 15, 408.

22. Vanderkraats,N.D., Hiken,J.F., Decker,K.F. and Edwards,J.R. (2013) Discovering high-resolution patterns of differential DNA methylation that correlate with gene expression changes. Nucleic Acids Res, 41, 6816–6827.

23. Fujiki,K., Shinoda,A., Kano,F., Sato,R., Shirahige,K. and Murata,M. (2013) PPARγ-induced PARylation promotes local DNA demethylation by production of 5-hydroxymethylcytosine. Nat Commun, 4, 2262.

24. Tuggle,K., Ali,M.W., Salazar,H. and Hooks,S.B. (2014) Regulator of G Protein Signaling Transcript Expression in Human Neural Progenitor Differentiation: R7 Subfamily Regulation by DNA Methylation. Neurosignals, 22, 43–51.

25. Huang,D.W., Sherman,B.T. and Lempicki,R.A. (2009) Systematic and integrative analysis of large gene lists using DAVID bioinformatics resources. Nat Protoc, 4, 44–57.

26. Spiers,H., Hannon,E., Schalkwyk,L.C., Smith,R., Wong,C.C.Y., O’Donovan,M.C., Bray,N.J. and Mill,J. (2015) Methylomic trajectories across human fetal brain development. Genome Res, 25, 338–352.

27. Weder,N., Zhang,H., Jensen,K., Yang,B.Z., Simen,A., Jackowski,A., Lipschitz,D., Douglas-Palumberi,H., Ge,M., Perepletchikova,F., et al. (2014) Child Abuse, Depression, and Methylation in Genes Involved With Stress, Neural Plasticity, and Brain Circuitry. Journal of the American Academy of Child & Adolescent Psychiatry, 53, 417–424.e5.

28. Schulz,H., Ruppert,A.-K., Herms,S., Wolf,C., Mirza-Schreiber,N., Stegle,O., Czamara,D., Forstner,A.J., Sivalingam,S., Schoch,S., et al. (2017) Genome-wide mapping of genetic determinants influencing DNA methylation and gene expression in human hippocampus. Nat Commun, 8, 1511.

29. Numata,S., Ye,T., Hyde,T.M., Guitart-Navarro,X., Tao,R., Wininger,M., Colantuoni,C., Weinberger,D.R., Kleinman,J.E. and Lipska,B.K. (2012) DNA Methylation Signatures in Development and Aging of the Human Prefrontal Cortex. The American Journal of Human Genetics, 90, 260–272.

30. Cecil,C.A.M., Walton,E., Smith,R.G., Viding,E., McCrory,E.J., Relton,C.L., Suderman,M., Pingault,J.-B., McArdle,W., Gaunt,T.R., et al. (2016) DNA methylation and substance-use risk: a prospective, genome-wide study spanning gestation to adolescence. Transl Psychiatry, 6, e976–e976.

31. la Rica,de,L., Deniz,Ö., Cheng,K.C.L., Todd,C.D., Cruz,C., Houseley,J. and Branco,M.R. (2016) TET-dependent regulation of retrotransposable elements in mouse embryonic stem cells. Genome Biol, 17, 234.

